# *Bacillus cereus* HS24 Suppresses Conidia Germination of *Magnaporthe oryzae* by Inhibiting the Ca^2+^ Signaling Pathway

**DOI:** 10.1101/310516

**Authors:** Wenxiang Huang, Xingyu Liu, Xiaosi Zhou, Yu Chen, Ye Li, Hongxia Liu

## Abstract

Rice yield is greatly reduced due to rice blast, a worldwide multi-cycle fungal disease caused by the ascomycete *Magnaporthe oryzae.* Previously, *Bacillus cereus* HS24 was isolated from rice growing area, which showed a strong bio-control effect on *M. oryzae.* In order to better exploit it as a bio-control agent, HS24 has been studied for its mechanism of controlling rice blast. Our results showed that conidial germination of *M. oryzae* was significantly inhibited by HS24 suspension, *n* and the inhibition rate reached to 97.83% at the concentration of 10 CFU/ml. The transcriptional level of *CAMKII, PMC1* and *CCH1,* which are key genes involved in Ca^2+^ signaling pathway, were significantly decreased in HS24-treated conidia. The treatment of *M. oryzae* with Ca^2+^ signaling pathway inhibitors KN-93, Verapamil, and cyclopiazonic acid (CPA) significantly reduced conidial germination rate and inhibited germ tube elongation. This inhibition effect was found to be concentration-dependent, similar to the HS24 treatment. By quantifying free Ca^2+^ in *M. oryzae* conidia, a significant reduction of intracellular free Ca^2+^ concentration in HS24-treated conidia in comparison to sterile water-treated conidia was found. The addition of exogenous Ca^2+^ did not abolish the inhibitory effect of HS24 on the reduction of intracellular free Ca^2+^ concentration and conidial germination. In conclusion, *B. cereus* HS24 can inhibit conidial germination by suppressing Ca^2+^ signaling in *M. oryzae,* and thus offers a great potential as a bio-control agent in rice blast management.

**Importance:** In the bio-control of rice blast, most of researches focused on the inhibitory effect of bio-control agents on development of *M. oryzae*, including inhibition of conidia germination, germ tube deformity, mycelium growth inhibition, etc, while little work has elucidated the molecular mechanisms underlying the various phenotypic change in *M. oryzae.* In order to better exploit HS24 as a potential bio-control agent, we studied the molecular mechanisms underlying the suppression of *B. cereus* HS24 on *M. oryzae* Guy11, and draw the conclusion that HS24 may inhibit conidia germination and germ tube elongation through inhibiting the Ca^2+^ signaling pathway. In this study, we characterized the morphological and physiological changes of *M. oryzae* when interacting with *B. cereus* HS24, and further investgated the responses of *M. oryzae* Ca^2+^ signallig pathway to HS24 treatment. Thus we have provided one more piece for the *B. cereus* / *M. oryzae* supression puzzle at the molecular level.

## Introduction

Rice *(Oryza sativa* L.) is the most important staple crop for over half of the world’s population. However, rice yield is greatly reduced due to rice blast, a worldwide multi-cycle fungal disease caused by the ascomycete *Magnaporthe oryzae (M. oryzae).* Infection by the pathogen is composed of a series of distinct processes (1). First, conidia adhere to the rice epidermis, and then germinate to form germ tubes under suitable environmental conditions. When the tips of germ tubes receive hydrophobic signals from the host, appressoria develop and invade the rice cells underneath. lnfectious hyphae ramify cellularly and intercellularly throughout host tissue thereafter. Visible symptoms become apparent 4-5 days after the initial adhesion. Secondary conidia are produced from the original lesion and spread to other parts of the host or other hosts (2). At present, rice blast management relies primarily on the utilization of resistant rice varieties and frequent chemical pesticide application. However, due to the limited choices of resistant rice varieties and the negative impact of chemical pesticides on environment (3), these two control methods present drawbacks. Additionally, fungicide resistant races emerge periodically because of selective pressure and the genetic complexity and diversity of *M. oryzae* (4). Biological control (bio-control) using antagonistic microorganisms has emerged as a promising, alterative control strategy, and has become the focus of extensive research in rice blast management. The action targets or function mode of microbial pesticides are generally multi-factory; therefore, there is less selective pressure on pathogens in comparison to conventional control methods (5). In addition, microbial pesticides are more environmentally sustainable than fungicide alternatives (6).

In recent decades, more and more bacteria have been found to be suppressive to *M. oryzae,* and some of them have been applied in agricultural production (5). So far, four sources of microbes have been developed into microbial pesticides against *M. oryzae,* namely *Bacillus, Pseudomonas,* fungi, and *Actinomycetes*, among which *Bacillus* is the most commonly and widely used (7). Yang and colleagues reported that a fresh culture of two new *B. etiophyticus* strains IPe2 and IPe14 had a bio-control efficacy of 64.4% and 56.4% respectively in reducing rice blast incidence (8). The water-soluble formulation of a *Bacillus megaterium* strain isolated in Thailand by Kanjanamaneesathian et al. has been reported to be effective in controlling rice blast and increasing yield (9). Saikia et al. isolated a *Brevibacillus laterosporus* strain BMP3 from hot spring soil in India, which achieved a 30% - 67% rice blast control efficacy under greenhouse conditions and prevented a yield loss of 35% - 56.5% (10). In a screening for rice blast bio-control agents, Ali et al. obtained multiple *Trichoderma* isolates from soil and demonstrated their disease control efficacy against *M. oryzae.* When used before transplanting rice seedlings, *Trichoderma* isolates alone or combined with *B. subtilis* UKM1 inhibited the occurrence of rice blast significantly (11).

With more bio-control agents being discovered, the mechanisms underlying their bio-control effects is more intriguing to researchers. A large number of studies have shown that *Bacillus* genus has an apparent antagonistic effect on *M. oryzae.* Suryadi and his colleagues found that *B. firmus* E65 had a significant inhibitory effect on mycelium growth of *M. oryzae* with an inhibition rate of 73% - 85% on solid culture media. *B. firmus* E65 was also working well in greenhouse, showing a bio-control efficacy of 50.4% against rice blast. Notably, panicle was best protected by *B. firmus* E65(12). Leelasuphakul et al. demonstrated the bio-control effect of *B. subtilis* NSRS89-24 culture filtrate on rice blast. The active antimicrobial substance purified from the culture filtrate was identified as dextranase, which can cause the cell wall to dissolve, and reduce the pathogenicity of *M. oryzae* (13). Tendulkar and colleagues purified a surfactant from *B. licheniformis* BC98 fermentation broth, which caused abnormal *M. oryzae* germ tube expansion, and further led to appressorium formation failure. This purified surfactant could also change the morphology of *M. oryzae* and inhibit the growth of mycelium (14). Zhang et al. found that *B. subtilis* KB-1122 could cause a series of deformities in *M. oryzae,* including germ tube abnormality, cell wall rupture and mycelium collapse. The serine, threonine, and tyrosine kinases are found to be up-regulated in KB-1122 when interacting with *M. oryzae,* suggesting that they may be involved in the mycelium collapse of *M. oryzae* (15). All these researches focused on the inhibitory effect of bio-control agents on development of *M. oryzae,* including inhibition of conidia germination, germ tube deformity, mycelium growth inhibition, mycelium collapse, etc, while little work has elucidated the molecular mechanisms underlying the various phenotypic change in *M. oryzae.* Conidia germination is the earliest essential step for growth, development and infection of *M. oryzae.* The Ca^2+^ signaling pathway is involved in the conidia germination of *yeast and filamentous fungal* (16, 17). The fluctuation of intracellular free Ca^2+^ concentration has an important regulatory effect on the metabolic activity of fungi. When the cells are subjected to environmental stimuli, the influx of extracellular Ca^2+^ and the release of intracellular Ca^2+^ reservior increase the free Ca^2+^ concentration in the cytoplasm, and the elevated Ca^2+^ concentration initiates the calcium signaling pathway (18). Key signaling factors Ca^2+^ channel proteins, Cch1, Ca^2+^ -ATPase Pmc1, and Ca^2+^/calmodulin-dependent protein kinase II (CaMKII), play an important role in the regulation of cellular Ca^2+^ concentration. The Ca^2+^ channel protein Cch1 is an important component of the high-affinity Ca^2+^ influx system, which localizes on the plasma membrane. When the high-affinity Ca^2+^ influx system is activated, it allows Ca^2+^ to flow from the extracellular space into the cell (18). Ca^2+^ -ATPase Pmc1 is a key protein localized on the vacuole membrane responsible for directing the cytosolic Ca^2+^ to vacuole (19). CaMKII is a serine/threonine-specific protein kinase and it is activated when the level of cytoplasmic Ca^2+^ is increased to regulate Ca^2+^ metabolism (20).

*Bacillus cereus* HS24 was isolated from rice rhizosphere in a field that was moderately contaminated with *M. oryzae.* HS24 demonstrated a bio-control efficiency of 77.5% in a 45 days greenhouse inoculation assay with *M. oryzae* Guy11 (21). In order to better exploit it as a potential bio-control agent, we studied the molecular mechanisms underlying the suppression of *B. cereus* HS24 on *M. oryzae* Guy11, and demonstrated that HS24 inhibits the conidia germination of *M. oryzae* by blocking its Ca^2+^ signaling pathway. This research has provided valuable information on how rhizobacteria *B. cereus* suppresses *M. oryzae,* and demonstrated its great potential as a bio-control agent in rice blast management.

## Materials and methods

### 1. Strains and culture conditions

*Activation and culture of HS24. B. cereus* HS24 was inoculated onto lysogeny broth (LB) agar plate (22), and incubated at 28 °C for 24 h in a constant temperature incubator. A single colony was picked and inoculated into 200 ml of liquid LB broth medium. The bacteria were cultured for 24 h under shaking conditions (200 rpm) at 28°C. HS24 fermentation broth was diluted to 1x10^5^, 1x10^6^, 1x10^7^ CFU/ml separately using sterile water. *Activation and culture of M. oryzae Guy11. M. oryzae* Guy11 (donated by Professor Zhengguang Zhang, Nanjing Agricultural University, Nanjing, Jiangsu, China) was inoculated onto potato dextrose agar (PDA) medium and cultured in a constant temperature incubator at 28 °C for 4 days in the dark for activation. Activated Guy11 was inoculated onto straw decoction and corn (SDC) agar media (23), and cultured at 28 °C in dark for 3-4 days, followed by 7 days of continuous illumination under fluorescent light to induce conidia production. 10 ml of sterile water was added to each plate, and a brush was used to collect conidia by gently brushing the surface of the plate. The conidia suspension was filtered through three layers of lens paper. The conidia was counted with a haemacytometer under microscope (24), and then the conida concentration was adjusted to 1 × 10^5^ conidia per milliliter using sterile water for use.

### 2. Screening of optimal HS24 working concentration

One milliliter of Guy11 conidia suspension was centrifuged in four 1.5 ml centrifuge tubes separately at 12000 rpm for 10 min to obtain conidia precipitates. 1 mL of sterile water as the control group and 1 mL of HS24 suspension with different concentrations (1x10^5^, 1x10^6^, 1x10^7^ CFU/ml) was added to each of the four centrifuge tubes to resuspend the conidia precipitates. Each treatment was repeated three times. Droplets (20 μl) of conidial suspension were placed on Fisher brand Microscope Cover Glass (hydrophobic) with wet filter paper and incubated at 28°C in darkness. Conidia germination, germ tubes extension and appressoria turgor pressure were monitored at different time points post incubation. Conidial germination and germ tube extension was examined under microscope (200X). More than 100 conidia were assessed for each treatment and each treatment had three repeats.

### 3. Effects of HS24 on appressoria turgor pressure and the germ tube length

The conidia of *M. oryzae* were treated with sterile water or HS24 at the concentration of 10^5^ CFU/ml, respectively. The germ tube length was measured at 6 hours post incubation (hpi) when it had stopped elongating. Glycerol solutions at various concentrations were used to detect the turgor pressure inside mature appressoria by cytorrhysis assay at 24 hpi (25).

### 4. RNA extraction and Ca^2+^ signaling involved genes transcriptional level detection

For real-time RT-PCR (RT-qPCR), *M. oryzae* conidia co-cultured with sterile water or *B. cereus* HS24 were collected after 2 hours of treatment for RNA extraction. After reverse transcription using a PrimeScript^TM^RT reagent Kit (TaKaRa, Code No. RR047a), the transcription of Ca^2+^ signaling involved genes was examined by qPCR. RT-qPCR reactions were performed following previously established procedures and each experiment was repeated three times (26). Q-PCR was performed with the ABI 7500 Fast real-time system (ABI Co.) and the transcripts were analyzed using the 7500 System SDS software (ABI Co.). The primers used in this study were designed on NCBI (http://www.ncbi.nlm.nih.gov) as shown in Table1.

### 5. Effect of Ca^2+^ signaling inhibitors on conidia germination and germ tube length

Ca^2+^ signaling inhibitors at different concentrations (EGTA, 0 mM / 50 mM / 100 mM / 150 mM / 200 mM; Verapamil, 0 μM / 100 μM / 200 μM / 300 μM / 400 μM; KN93, 0 μM / 20 μM / 40 μM / 60 μM / 80 μM / 100 μM; CPA, 0 μM / 20 μM / 40 μM / 60 μM / 80 μM / 100 μM) were added to the conidia suspension. 20 μl of the mixture was dripped onto Fisher brand Microscope Cover Glass (hydrophobic) with wet filter paper and incubated at 28°C in darkness. After 6 h of incubation, conidia germination rate and germ tube length were detected.

### 6. Determination of Ca^2+^ concentration in conidia

The free Ca^2+^ in the conidia of *M. oryzae* was stained with Fluo 3-AM (Sigma, Code No. 39294) at the concentration of 150 μM, referring to the method of Zhang et al (27). The change of Ca^2+^ concentration was indicated by relative fluorescence intensity. Relative fluorescence intensity = the fluorescence intensity of each treatment / the fluorescence intensity of water treatment.

### 7. Statistics and Data analysis

All treatment groups were repeated at least three times and the data was analyzed with SPSS 17.0 by LSD (least significant difference test) analysis.

## Results

### 1. *B. cereus* HS24 at the concentration of 1×10 CFU/ml inhibited the conidia germination

Conidia of *M. oryzae* were treated with various concentrations of HS24 suspension. Conidia germination rate was examined at different time points and the morphological changes of germ tubes and appressoria were monitored. The results showed that at 8 hpi the conidia germination rate of the sterile water-treated group reached to 93%, while the germination rate of the three HS24-treated groups (1×10^5^, 1×10^6^, 1×10^7^ CFU/ml) were 45.04%, 17.5%, and 3% respectively. After 12 hours of co-culture, the conidia germination rate of the HS24-treated groups (1×10^5^, 1×10^6^, 1×10^7^ CFU/ml) were 49.13%, 60.25%, 95%, respectively). At 24hpi the germination rate of the 1x10 CFU/ml HS24-treated group was 2.17% (Fig. 1A). The above results showed that HS24 at the concentration of 1x10^7^ CFU/ml was able to significantly inhibit the conidia germination of *M. oryzae.* The inhibitory effect on conidia germination became weaker as the HS24 concentration decreased (Fig. 1A). No significant difference on conidia germination rate was found between the treatments of sterile water and LB broth, so we used the sterile water as the control group in the rest of the experiments. Although the germ tube length of conidia treated with HS24 at the concentration of 1x10^5^ CFU/ml was significantly shortened, circular melanin-containing mature appressoria were abundantly observed (Fig. 2). In addition, no significant changes of turgor pressure was observed between water or HS24 co-cultured *M. oryzae* samples when the appressoria were treated with glycerol at various concentrations (Fig. 3). The results showed that *B. cereus* HS24 at the concentration of 1x10^5^ CFU/ml was able to significantly inhibit the conidia germ tube elongation, but had not significant effect on the formation and turgor pressure of appressoria.

**Fig. 1.**
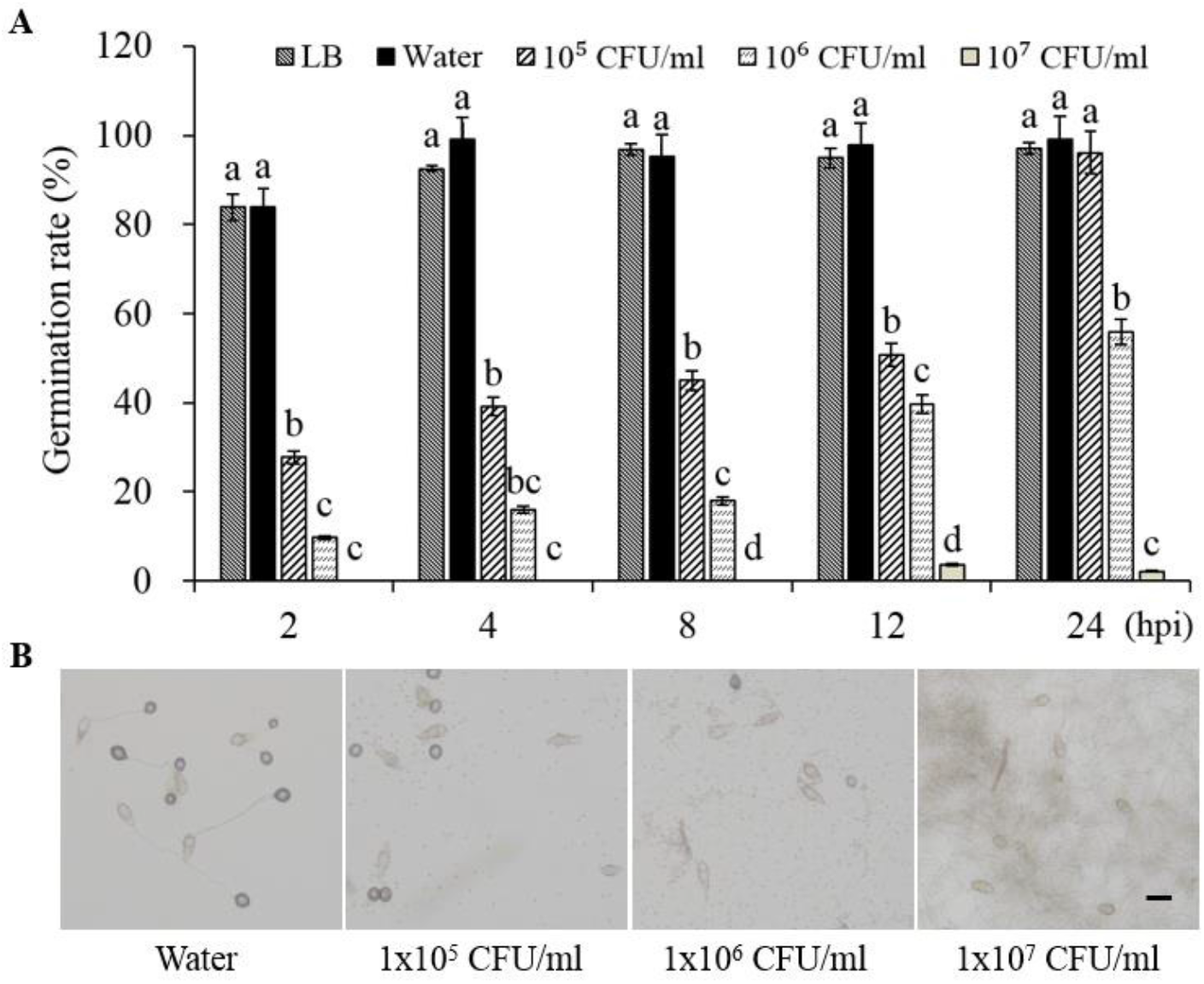
Effect of HS24 at various concentrations on the conidia germination rate of*M. oryzae* The Conidia of *M. oryzae* were treated with LB broth, sterile water (Control) and bacterial suspension (HS24) at the concentration of 1x10^5^, 1x10^6^, 1x10^7^ CFU/ml. (A) Germination rate of *M. oryzae* conidia was evaluated at 2, 4, 8, 12 and 24 h post incubation with sterile water or HS24 suspension. Statistically significant difference (P < 0.05) was indicated by bars with different letters. (B) Conidia of *M. oryzae* were examined under a microscope at 24 h post incubation with sterile water or HS24 suspension, bar = 20 μm.

**Fig. 2.**
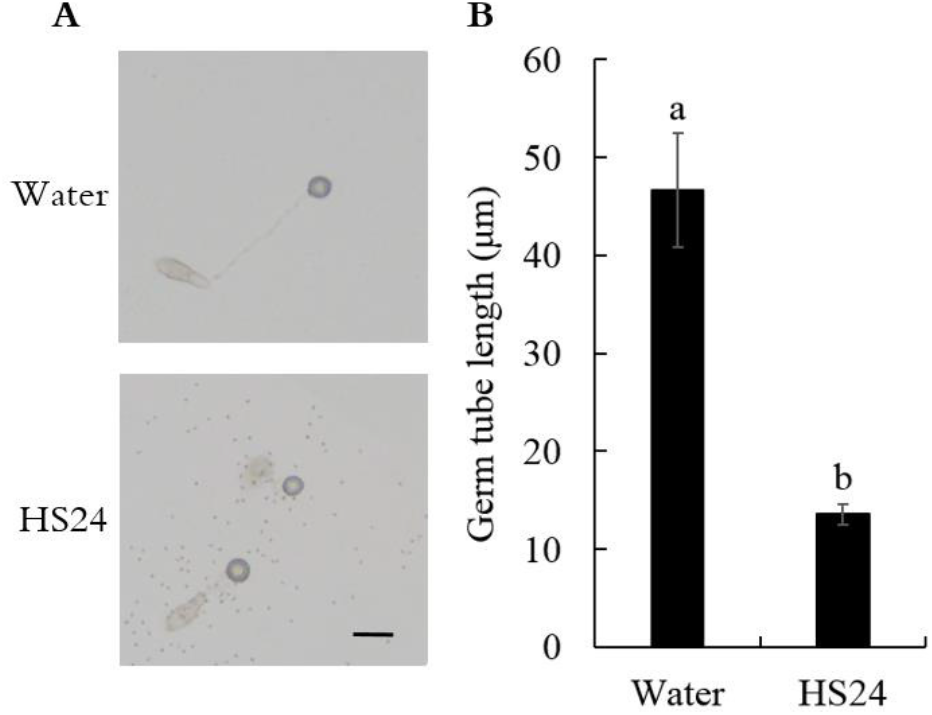
Effect of HS24 on germ tube length at the concentration of 1 × 10^5^ CFU/ml (A) Conidia were treated with sterile water (Control) or bacterial suspension (HS24) at the concentration of 1×10^5^ CFU/ml. Bar = 20 μm. (B) The germ tube length was measured at 6 h post incubation. Statistically significant difference (P < 0.05) was indicated by bars with the different letters.

**Fig. 3.**
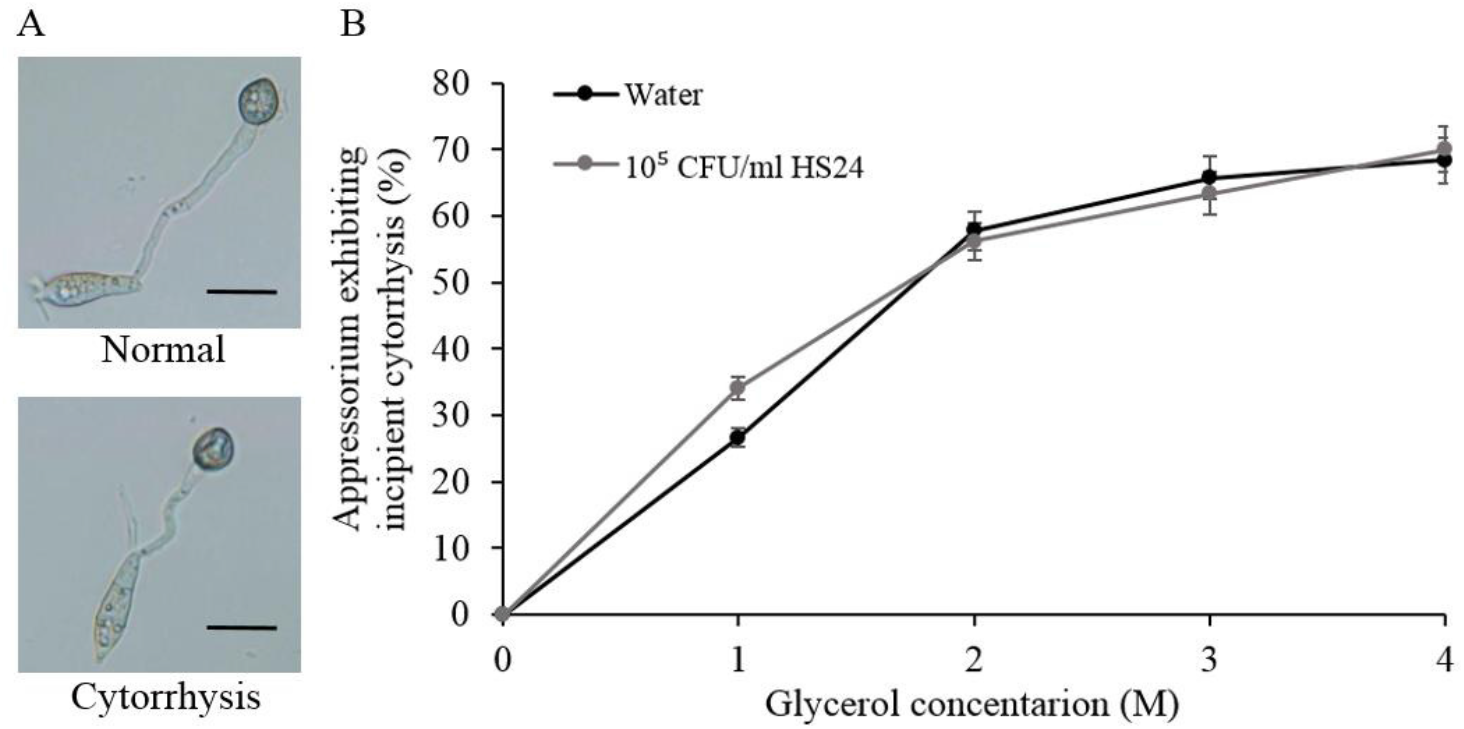
The effect of HS24 on turgor pressure in appressoria at the concentration of 1×10^5^ CFU/ml (A) Normal and cytorrhysis appressoria were observed from both water-treated or HS24-treated samples. Upper panel, normal appressorium; Lower panel, cytorrhysis appressorium. Bar=20 μm. (B) Cytorrhysis rate of appressoria treated with glycerol at various concentrations 24 hpi.

### 2. HS24 down-regulated the transcriptional level of genes involved in *M.oryzae* Ca^2+^ signaling pathway

In this study, transcription level of three key Ca^2+^ signaling involved genes, Ca^2+^/calmodulin-dependent protein kinase II coding gene *CaMKII*, Ca^2+^-ATPase coding gene *PMC1,* and high-affinity Ca^2+^ influx system protein coding gene *CCH1* were detected by RT-qPCR. The result showed that the transcriptional level of these three genes in HS24-treated conidia were significantly down-regulated (Fig. 4).

**Fig. 4.**
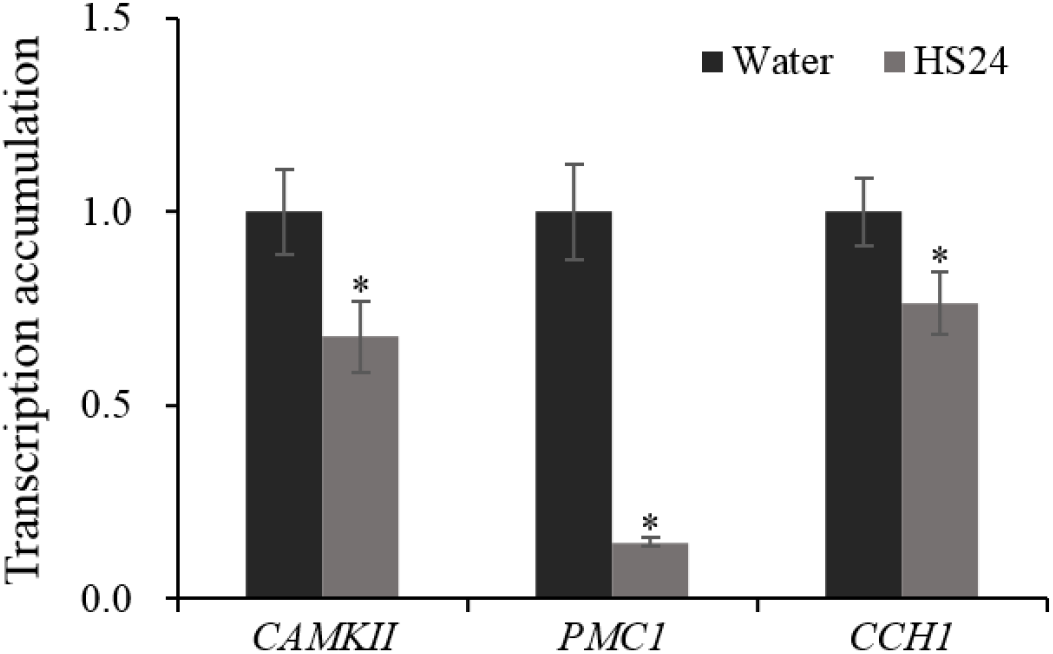
The Ca^2+^ signaling involved genes transcription accumulation in HS24-treated conidia *M. oryzae* conidia were treated with sterile water or bacterial suspention (1×10^7^ CFU/ml HS24) for two hours and then the total RNA was extracted. RT-qPCR detection of gene transcription accumulation was carried out after reverse transcription of mRNA. Small asterisk represents a significant difference at 0.05 level (P < 0.05).

### 3. Ca signaling inhibitors inhibited conidia germination and germ tube elongation

It has been observed that the transcriptional level of *CaMKII*, *PMC1* and *CCH1* were significantly down-regulated in condia treated by HS24. In order to test whether the down-regulation of these signaling factors led to the inhibition of conidia germination and germ tube elongation, the conidia of *M. oryzae* were treated with corresponding signaling inhibitors. The results showed that Ca^2+^ chelating agent EGTA, CaMKII inhibitor KN-93, calcium channel protein Cch1 inhibitor Verapamil, and Ca^2+^ -ATPase Pmc1 inhibitor CPA all inhibited the conidia germination, and the inhibitory effect increased with the inhibitiors concentration increased (Fig. 5A, C, E, G). This concentration-dependent mannerwas the same as HS24 treatment. Meanwhile, the length of germ tube was also affected by the inhibitors (Fig. 5B, D, F, H). These experimental results further demonstrated that HS24 inhibits the conidia from germination by inhibiting gene expression of key elements in Ca^2+^ signaling pathway.

**Fig. 5.**
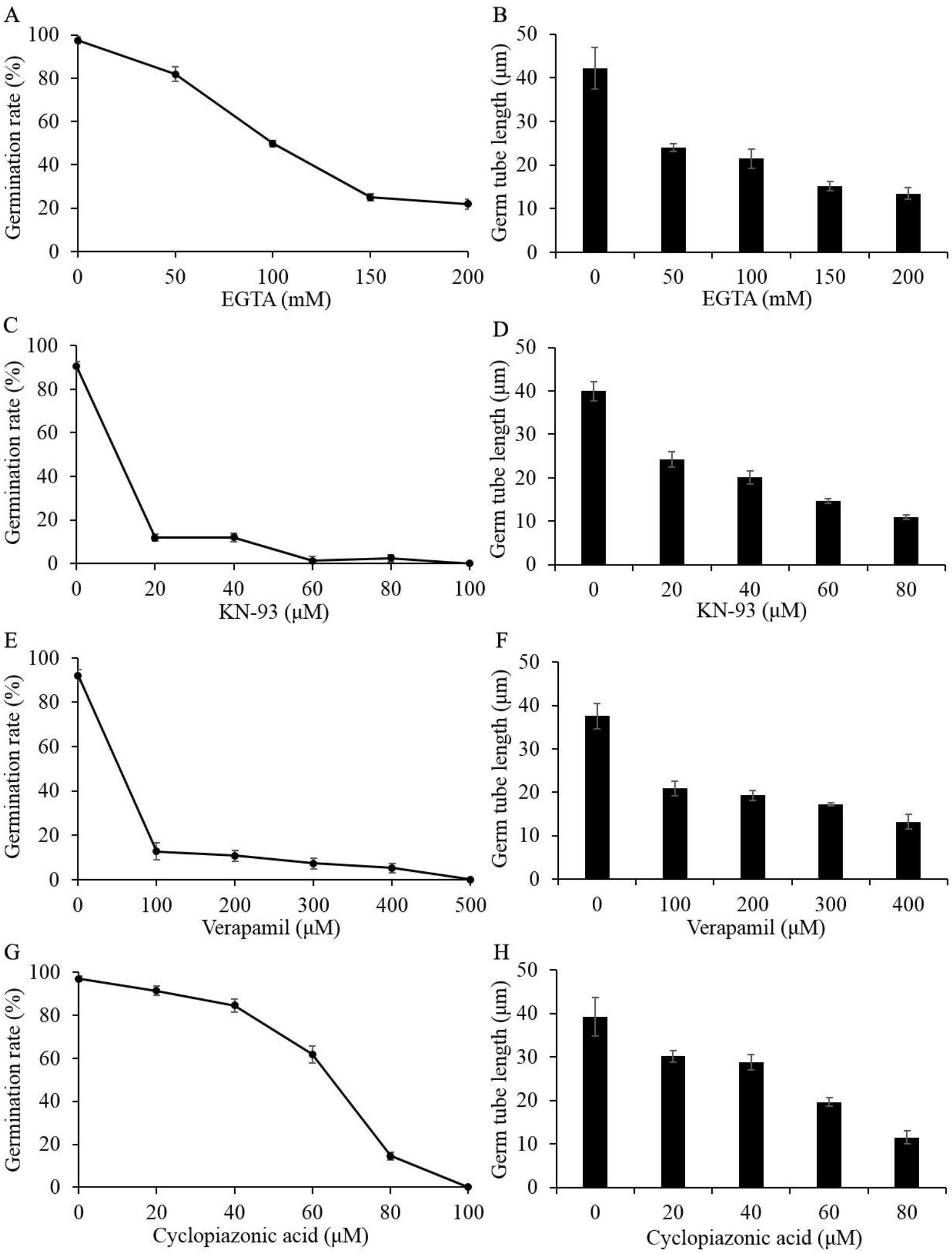
Effect of exogenous addition of EGTA, KN-93, Verapamil and cyclopiazonic acid on conidia germination rate and germ tube length of Guy11. (A), (C), (E), (G) Effect of various concentrations of inhibitors on the conidia germination rate. (B), (D), (F), (H) Effect of various concentrations of inhibitors on germ tube length of*M. oryzae* conidia.

### 4. HS24 treatment led to the appearance of Granular vesicles in *M. oryzae* conidia

It has been reported that the treatment of Ca^2+^ signal inhibitors Verapamil and KN-93 resulted in the appearance of granular vesicles in the uniform protoplasm of *M. oryzae* conidia (16). In order to test whether HS24 has a similar effect, *M. oryzae* conidia were treated with HS24 suspension. Granular vesicles were found in the HS24-treated conidia but not in the water-treated conidia (Fig. 6A). This result further illustrated that HS24 suppresses *M. oryzae* conidia germination through interfering its Ca^2+^ signaling pathway.

**Figure 6.**
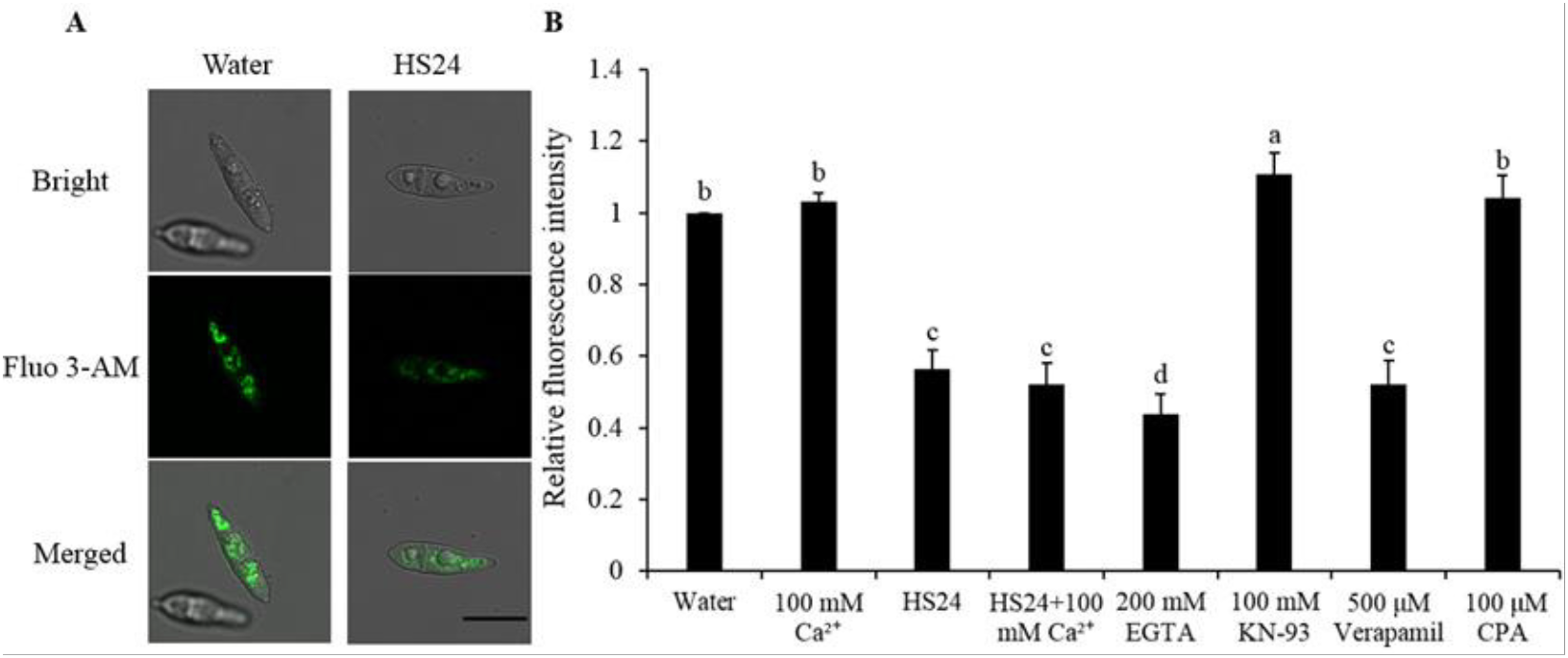
Free Ca^2+^ concentration and appearance of granular vesicles within the conidia treated with HS24 (A) Ca^2+^ fluorescence staining of conidia treated by sterile water or bacterial suspension (1×10^7^ CFU/ml HS24). (B) The fluorescence intensity of free Ca^2+^ in conidia. Conidia with various treatments were incubating with fluorescent dye Fluo 3-AM for 2 hours at 4 °C. Statistically significant difference (P < 0.05) was indicated by bars with the different letters.

### 5. HS24 reduces the intracellular Ca^2+^ concentration

Since HS24 inhibits the expression of key genes involved in Ca^2+^ signaling, we would like to further test whether HS24 treatment could change intracellular free Ca^2+^ concentration in conidia. Free intracellular Ca^2+^ was stained with Ca^2+^ fluorescent dye Fluo 3-AM, and free Ca^2+^ concentration was indicated by relative fluorescence intensity. As the result shown, HS24, EGTA and verapamil-treated conidia showed weaker fluorescence intensity compared to the sterile water treatment (Fig. 6A), and the added exogenous Ca^2+^, KN-93 and CPA increased the concentration of free Ca^2+^ in the conidia. These experimental results indicate that HS24 can interfer the Ca^2+^ signaling pathway by abolishing the function of CaMKII, Pmc1 and Cch1 and thus result in the inhibition of conidia germination and germ tube elongation. Exogenous addition of various concentrations of calcium ions under HS24 treatment showed that the addition of exogenous Ca^2+^ failed to compensate the inhibitory effect of HS24 on conidial germination (Fig. 7).

**Fig. 7.**
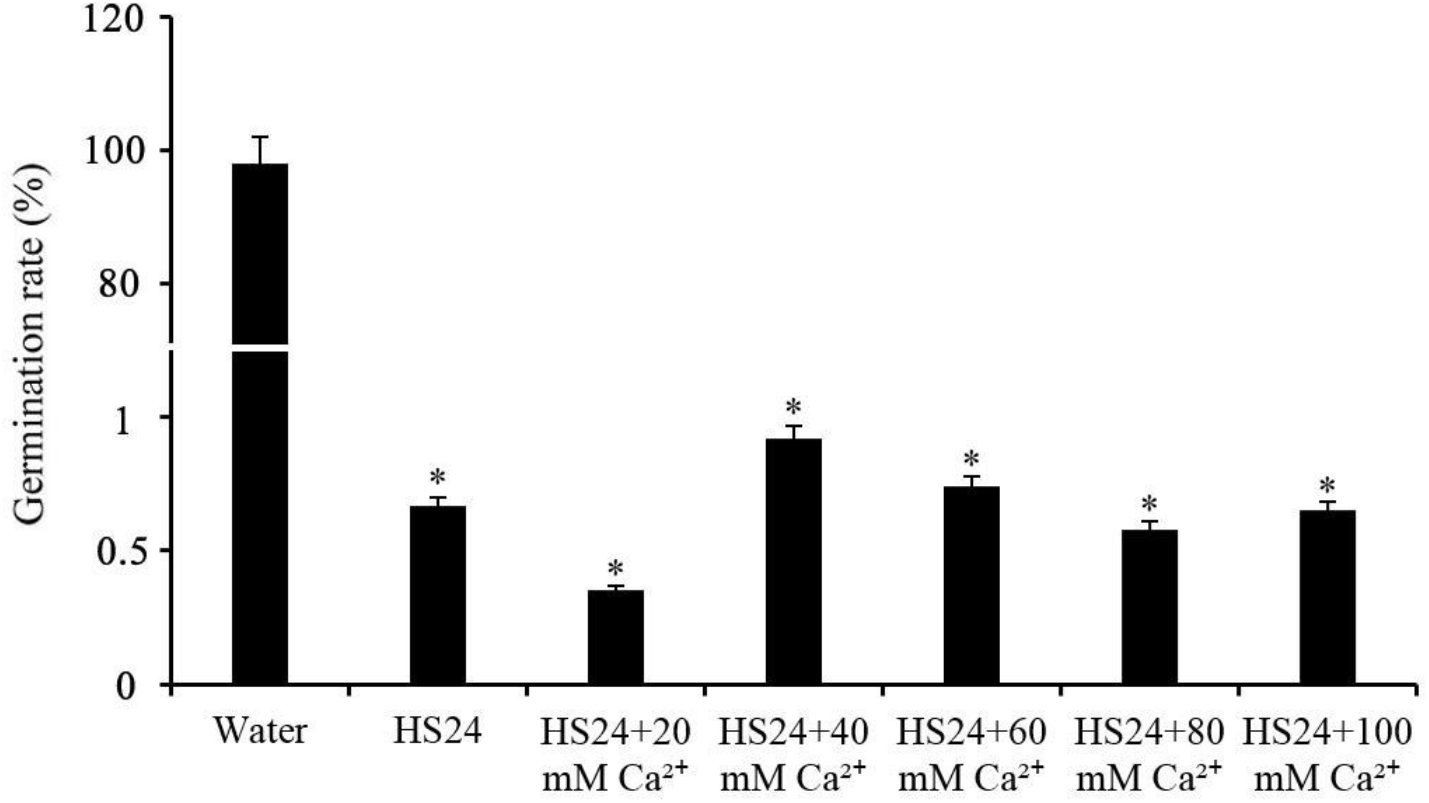
Influence of exogenous Ca^2^on the conidia germination rate treated by HS24 HS24 concentration was 1 × 10^7^ CFU/ml. Small asterisk represents a significant difference at 0.05 level compared with the sterile water treatment (P < 0.05).

## Discussion

In this research, we mainly studied the effect of *B. cereus* HS24 on *M. oryzae* with an aim to better understand the mechanism of HS24 as an agent for the bio-control of rice blast. The results show that HS24 has a strong antimicrobial effect on *M. oryzae* through interfering with the Ca^2+^ signalling pathway to inhibit conidia germination and germ tube elongation. *Bacillus* genus is a famous antiagnistic bio-control angent against pathogenic fungi and the bio-control mechanism mainly involves antagonism, in which the inhibition of mycelia growth and conidia germination is largely reported. Shan and associates isolated *B. methylotrophicus* BC79 from Shanxi Qinling and found that BC79 culture filtrate had a bio-control efficacy of 89.9% against *M. oryzae* under greenhouse conditions. They further identified phenaminomethy-lacetic acid as the major inhibitory substance produced by BC79, which plays a key role in inhibition. The purified phenaminomethylacetic acid can completely inhibit the blast *M. oryzae* conidia germination, and lead to hyphae deformity and even cause cell death (28). In a screening for natural alternatives to synthetic fungicides to control rice blast, Li et al. found that citral has a strong antifungal effect against *M. oryzae.* Citral was able to inhibit conidia germination, suppress germ tube elongation, restrain mycelium growth, and prevent colony development in a concentration-dependent manner *in vitro.* After exposure to citral, the surface of mycelium cells observed under a scanning electron microscope (SEM) became wrinkled, the mycelium cell wall became thin, the cell membran was impaired, and the villus-like material outside the cell wall was lost (29). Sha et al. studied the interactions between two *B. subtilis* strains SYX04 and SYX20 and *M. oryzae.* They found that SYX04 and SYX20 not only significantly inhibited *M. oryzae* conidia germination, germ tube elongation and attachment formation, but also caused series of structural changes on hypha and conidia. Observation with SEM and transmission electron microscopy (TEM) revealed that the cell wall and membrane structure of *M.oryzae* showed ultrastructural abnormalities with severe degradation (30). These studies have reported the isolation of biological control strains, or the identification of active substances that have the effect of inhibiting conidia germination and germ tube elongation. However, these studies focused on observation of the morphological changes of *M. oryzae,* and little work has illucidated the molecular mechanisms causing these changes.

In order to reveal the molecular mechanisms of the suppressive effect of HS24 on *M. oryzae,* we studied the influence of HS24 on *M. oryzae* Ca^2+^ signalling pathway. Intracellular Ca^2+^ is an important second messenger in all organisms. When the Ca^2+^ concentration in the environment varies from 1 mM to 100 mM, there is a slight change in the concentration of Ca^2+^ inside the biological cytoplasm, usually in the range of 50 nM to 200 nM (31–33). The calcium homeostasis system, composed of various calcium channels and ion pumps, as well as many related proteins and enzymes, plays an important role in maintaining optimum Ca^2+^ concentrations in the cytoplasm and in organelles such as vacuoles, endoplasmic reticulum, and Golgi apparatus (19, 31, 34). Normally, the plasma membrane Ca^2+^ influx system is activated in response to various external environmental changes such as alkaline stress, cold / heat stress, oxidative stress and ethanol stress (35–39). The transient increase in intracellular concentration can also be induced by Ca^2+^ secretion from the internal calcium pool (18). The increased Ca^2+^ concentration in yeast and filamentous fungal cells directly affects a wide range of cellular processes such as sporulation, conidia germination, mycelium tip growth, mycelium branching, gene expression, and circadian rhythms. This increased Ca^2+^ concentration also regulates the signaling cascade and activates the calcineurin pathway to reduce Ca^2+^ concentration to basal level (35, 40). In this study, the transcriptional level of Ca^2+^/calmodulin-dependent protein kinase II coding gene *CAMKII,* Ca^2+^ -ATPase coding gene *PMC1* and Ca^2+^ channel protein coding gene *CCH1* were down regulated in conidia of *M.oryzae* treated with HS24, and we hypothesized that the founction of the corresponding protiens were affected to some extent. The same experimental results were obtained by exogenous addition of Ca^2+^ signaling inhibitors. Both treatments caused inhibition of conidia germination at high concentration and inhibition of germ tube elongation at low concentration. The decreased *M. oryzae* conidia germination rate cuased by CaMKII inhibitor KN-93 was consistent with the decreased *Colletotrichum gloeosporioides* conidia germination rate treated with KN-93 (41). The inhibitory effect of EGTA and Cch1 inhibitor Verapamil on reducing *M. oryzae* conidia germination rate was supported by the experimental results of Wang et al. They also reported that Verapamil and KN-93 caused the appearance of granular vesicles in uniform protoplasm within the conidia (16), and the same phenomenon occurred when the conidia were treated with HS24 (Fig. 6A), which further demonstrated the inhibition of HS24 on the Ca^2+^ signaling pathway of *M. oryzae.* The chelation of exogenously added EGTA with free Ca^2+^ resulted in a dramatic decrease of the calcium concentration in environment, and caused the decrease of conidia germination rate (Fig. 5A), but the inhibition of conidia germination by HS24 was not eliminated by the addition of exogenous Ca^2+^ (Fig. 7). This result suggested that HS24 could cause a decrease of free Ca^2+^ concentration in the conidia by preventing free Ca^2+^ entering the conidia from the environment. Free Ca^2+^ labeling with fluorescent dye in HS24-treated conidia revealed a significant decrease of intracellular free Ca^2+^ concentration, accompanied by the down-regulation of transcription levels of three key Ca^2+^ signaling molecules. Taken the above results together, we draw the conclusion that HS24 may inhibit conidia germination and germ tube elongation through the following pathways. First, HS24 blocks extracellular Ca^2+^ influx by inactivating high-affinity Ca^2+^ influx system protein Cch1 (Fig. 4, Fig. 6) and the reduced Ca^2+^ concentration inhibits calcium signaling pathway. Second, HS24 affects the activity of other key factors of the calcium signal pathway such as CaMKII and Pmc1 (Fig. 4, Fig. 6), and thus interfer Ca^2+^ signaling. In this study, we characterized the morphological and physiological changes of *M. oryzae* when interacting with *B. cereus* HS24, and further investgated the responses of *M. oryzae* Ca^2+^ signallig pathway to HS24 treatment. Thus we have provided one more piece for the *B. cereus/M. oryzae* supression puzzle at the molecular level.

**Table 1.**
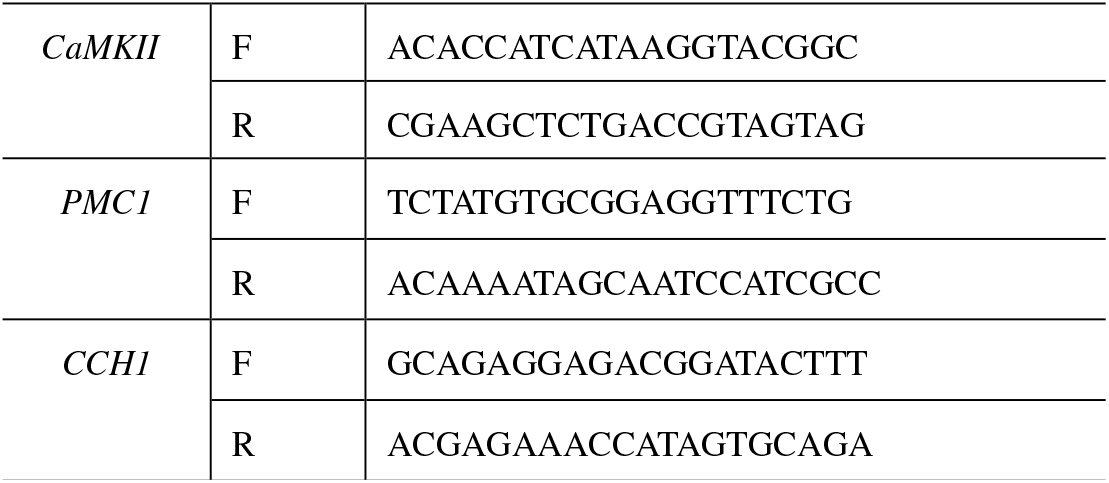
The primers used in this study

## Acknowledgements

This work was supported by National Natural Science Foundation of China (31571992, 31371925). We greatly appreciate professor Zhang, College of Plant Protection, Nanjing Agricultural University, for the donation of Guy11 strain.

